# Automated Bacteriophage Evolution Studies with the Aionostat

**DOI:** 10.1101/2025.04.16.649189

**Authors:** Valentin Druelle, Marco Molari, Giacomo Castagnetti, Alexander Harms, Richard A. Neher

## Abstract

Bacteriophages, the viruses that infect bacteria, are the most abundant and diverse biological entities on our planet. They play a critical role in shaping ecosystems and are increasingly recognized for their potential in treating bacterial infections. Yet, our comprehension of their biology and evolutionary dynamics is limited, largely because research has concentrated on a select few well-characterized phages or relies on broad metagenomic studies with limited follow-up analysis of individual phages. This knowledge gap hinders our capacity to exploit their therapeutic and ecological possibilities – and while some studies have attempted to bridge it, such efforts typically require a lot of manual labor, highlighting the need for high-throughput, reproducible methods for in-depth study of phage evolution.

To address this gap, we introduce the Aionostat, a novel automated continuous culture device designed to facilitate bacteriophage directed evolution experiments at scale. The Aionostat’s potential is showcased through two example experiments. In the first, phages from the BASEL collection rapidly adapted to a challenging *E. coli* strain, acquiring mutations and deletions that improved their infectivity. In the second experiment, we evolved a mixture of these phages on the same *E.coli* strain, leading to the emergence of recombinant phages with increased fitness. By automating these experiments, the Aionostat enables faster, more reproducible studies of phage evolution that would be impractical to perform by hand, thereby opening new avenues for investigating viral dynamics, engineering phage therapies, and studying evolutionary principles in broader biological contexts.

## Introduction

Bacteriophages are of significant biological interest due to their role in shaping microbial ecosystems thanks to their extensive diversity (Dion et al., 2020; Parmar et al., 2018; Chevallereau et al., 2022; Mahmoudabadi, Phillips, 2018). Their potential as therapeutic agents, especially against antibiotic-resistant bacteria, has been demonstrated in multiple studies (Gordillo Altamirano, Barr, 2019; Schmidt, 2019; Doss et al., 2017; Caflisch et al., 2019). For effective therapeutic outcomes, bacteriophages must often be specifically adapted to target bacteria (Abdelsattar et al., 2021; Borin et al., 2021). This adaptation is typically achieved using directed evolution experiments. A popular protocol, known as Appleman’s protocol, leverages the natural ability of bacteriophages to evolve via both vertical inheritance and horizontal gene transfer by engaging in recombination with other phages (Burrowes et al., 2019; Hendrix, 2002; Hatfull, Hendrix, 2011; Mavrich, Hatfull, 2017). Directed evolution allowing for recombination of a diverse genetic pool reliable produces phages with the ability to infect challenging bacterial strains (Loose et al., 2021). However, the role of experimental parameters and evolutionary mechanisms to optimize phages are not well understood, emphasizing the importance of understanding phage evolution (Ofir, Sorek, 2018). Beyond their direct environmental and therapeutic applications, bacteriophages also serve as a valuable model for viral evolution and can be linked to broader studies of protein and RNA evolution (Turgeon et al., 2014; Brödel et al., 2018). Techniques such as phage display exemplify how attributes tied to phage fitness can be enhanced through the evolutionary processes of the phages themselves (Willats, 2002).

The current state of phage evolution research is limited to a handful of well characterized bacteriophages (Keen, 2015), or broad environmental metagenomics phage studies where the phages themselves are rarely isolated and studied individually (Sousa et al., 2020; Bonilla-Rosso et al., 2020; Tisza, Buck, 2021). The former limits the scope of the findings, while the latter prevents detailed analysis that would require experimental intervention. To better understand and manipulate phage evolution to our benefit we need to bridge this gap, which requires methods that are high-throughput, rapid, reproducible, and cost-effective.

To allow for more systematic investigations of phage evolution at higher throughput, we developed the Aionostat: an automated continuous culture device designed for directed evolution experiments of bacteriophages. The name Aionostat is derived from its similarity to other devices such as chemostats and turbidostats, and as a reference to the Greek deity Aion who symbolizes cyclical time (Novick, Szilard, 1950; Toprak et al., 2012). This reflects the device’s continuous and iterative approach to cultivating and evolving phages. The device operates similarly to a morbidostat, with some modifications that improve performance, ease of use, reliability and make it phage-compatible (Toprak et al., 2013). It can be seen as improvement of similar phage continuous culture devices thanks to its versatility (Holtzman et al., 2020; Brödel et al., 2020). The most basic setup used for phage evolution is a dual-vial system as shown in Figure 1.A. The first vial maintains bacteria in their exponential growth phase without the presence of phages. These bacteria are then channeled to the second vial where they encounter and get infected by phages. Over the replications cycles, genomic changes can appear in the newly produced virions. Beneficial mutations confer a selective advantage, leading to their predominance within the phage population, ultimately resulting in phages with superior bacterial adaptation as illustrated in Figure 1.C.

**Figure 1:**
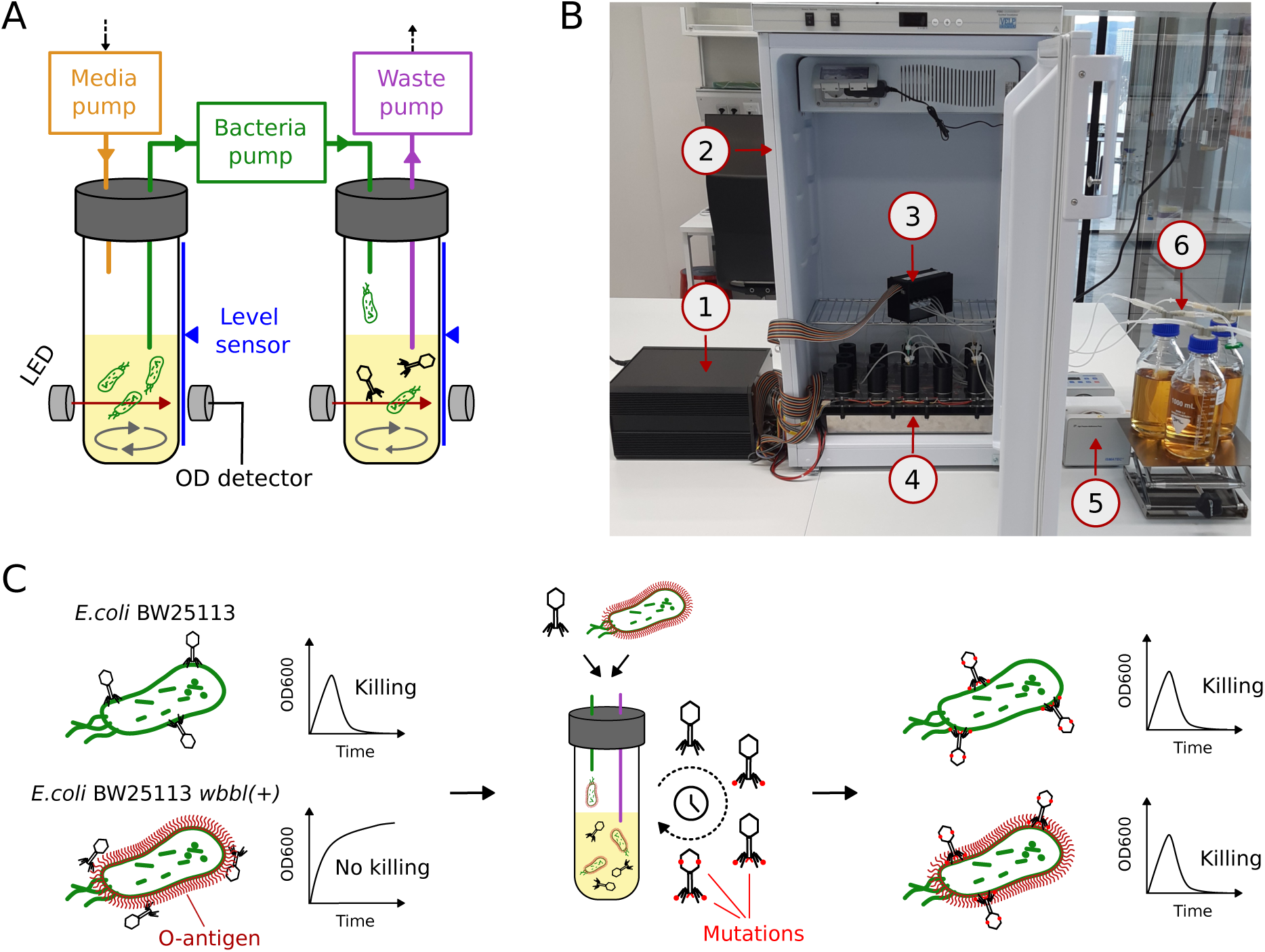
Introducing the Aionostat. (**A**) Schematic of the working principle of the Aionostat. Fresh media is pumped in the bacterial growth vial, where a culture of bacteria is maintained in exponential phase. This provides fresh bacteria to the phage vial, where phages are grown and evolved. (**B**) Picture of the device. 1: electronic components’ enclosure 2: incubator for temperature control 3: single channel pump array 4: array of experiment vials on stirrer plate 5: peristaltic exhaust pump 6: input media bottle. The waste bottle is not shown on the picture. (**C**) Schematic of the evolution experiment. Originally the phage does not infect the strain with restored O-antigen well, resulting in limited bacterial killing when the bacteria are grown in presence of the phage. After evolution using the Aionostat, the evolved phages infect both the ancestor and the O-antigen strain, causing significant bacterial killing.

In this paper we present two phage evolution experiments performed with the Aionostat to showcase its potential. First, a ‘linear’ evolution experiment where *E.coli* bacteriophages from the BASEL phage collection (Maffei et al., 2021) were selected and evolved in separate vials for better infectivity on a challenging (decorated with O-antigens) *E.coli* strain. The fitness of the evolved phages was measured and compared to their ancestors. Second, a phage “cocktail” experiment where these phages were mixed and evolved on the same challenging *E.coli* strain, which resulted in the appearance of recombinant phages.

The Aionostat was designed to meet the research challenges previously outlined, offering a methodological improvement over existing systems and enabling high throughput and reproducible evolution phage experiments that would otherwise be too labor-intensive to perform by hand. Furthermore, its versatility makes it adaptable to various applications beyond those described, providing an enriched platform for bacteriophage and bacterial research. Below, we describe two experiments conducted with the Aionostat and demonstrate how it can be used to systematically investigate phage evolution at the level of individual mutations and complex recombination patterns between co-cocultured phages. We provide details about the Aionostat, how it is built, how it works and how to use it in the Materials and Methods section and in Supplementary Figure 1.

## Results

In this study we present the ability of the Aionostat to perform directed phage evolution to enhance their fitness on a challenging bacterial strain that they do not infect well originally. This was done in two experiments, a linear evolution experiment and a phage “cocktail” experiment which we present in the following sections.

We chose three bacteriophages belonging to the *Vequintavirinae* subfamily and its relatives as phage models for our directed evolution experiments. These bacteriophages are designated as phage WalterGehring (Bas51, NCBI GenBank accession MZ501111.1), phage MaxBurger (Bas54, accession MZ501093.1), and phage PaulScherrer (Bas60, accession MZ501100.1), and we refer to them via their Bas identifier. They are all lytic bacteriophages. The phages were sourced from the BASEL phage collection as it is a set of well characterized phages (Maffei et al., 2021). We chose this group not only because they grow well, but also due to their comparably large, mosaic genomes – indicating their ability to recombine – and because they were able to evolve the phenotype wanted. The *Vequintavirinae* family (and relatives) is well represented in this collection, providing a robust foundation for our research. These phages have a remarkably large tail fiber locus, with multiple co-expressed tail fibers and tail spikes. This is a peculiar but shared feature among this group of phages, which likely enables them to attach to a variety of surface motifs on the bacterial surface. We assess the directed evolution of the three phages against a challenging strain, *E.coli* BW25113 *wbbl(+)* (Maffei et al., 2021). This strain is a variant of the commonly used *E.coli* K-12 BW25113 strain in which the O16-type O-antigen glycan barrier was restored (Liu, Reeves, 1994). It is otherwise genetically similar to the isolation strain of the phages. The restored O-antigen adds long chains on the lipopolysaccharide (LPS), which act as a protective barrier on the bacterial surface by shielding the cell surface as shown in Figure 1.C. This inhibits infection from bacteriophages that do not bind the LPS or other glycans (Maffei et al., 2021). The phages used in this study are impaired by the O-antigen, but infection is not completely inhibited. This is likely due to their ability to bind another surface glycan, which apparently enables them to bypass the O-antigen barrier (Sellner et al., 2021).

### The Aionostat enables automation of phage evolution experiments

The experiments were conducted using the Aionostat over a duration of 10 days, as depicted in schematic Figure 2.A and Figure 3.A. Phage vials are paired with a culture vial of *E.coli* BW25113 *wbbl(+)* kept in exponential phase in lysogeny broth medium (LB) at 37°C. Vials were seeded from bacterial and phage stocks right before the start of the experiment. The bacterial culture’s dilution rate with raw LB was automatically adjusted to maintain a constant optical density at 600nm (OD600) of 0.4. The excess liquid resulting from the bacterial culture dilution was transferred to the phage vial, where infection and replication of the phages occur. This transfer of bacterial culture to the phage vial provides fresh bacteria for infection, while the volume in the vial is maintained by discarding any surplus. Consequently, the phage solution gets diluted over time, which selects for bacteriophages with higher fitness on *E.coli* BW25113 *wbbl(+)*.

**Figure 2:**
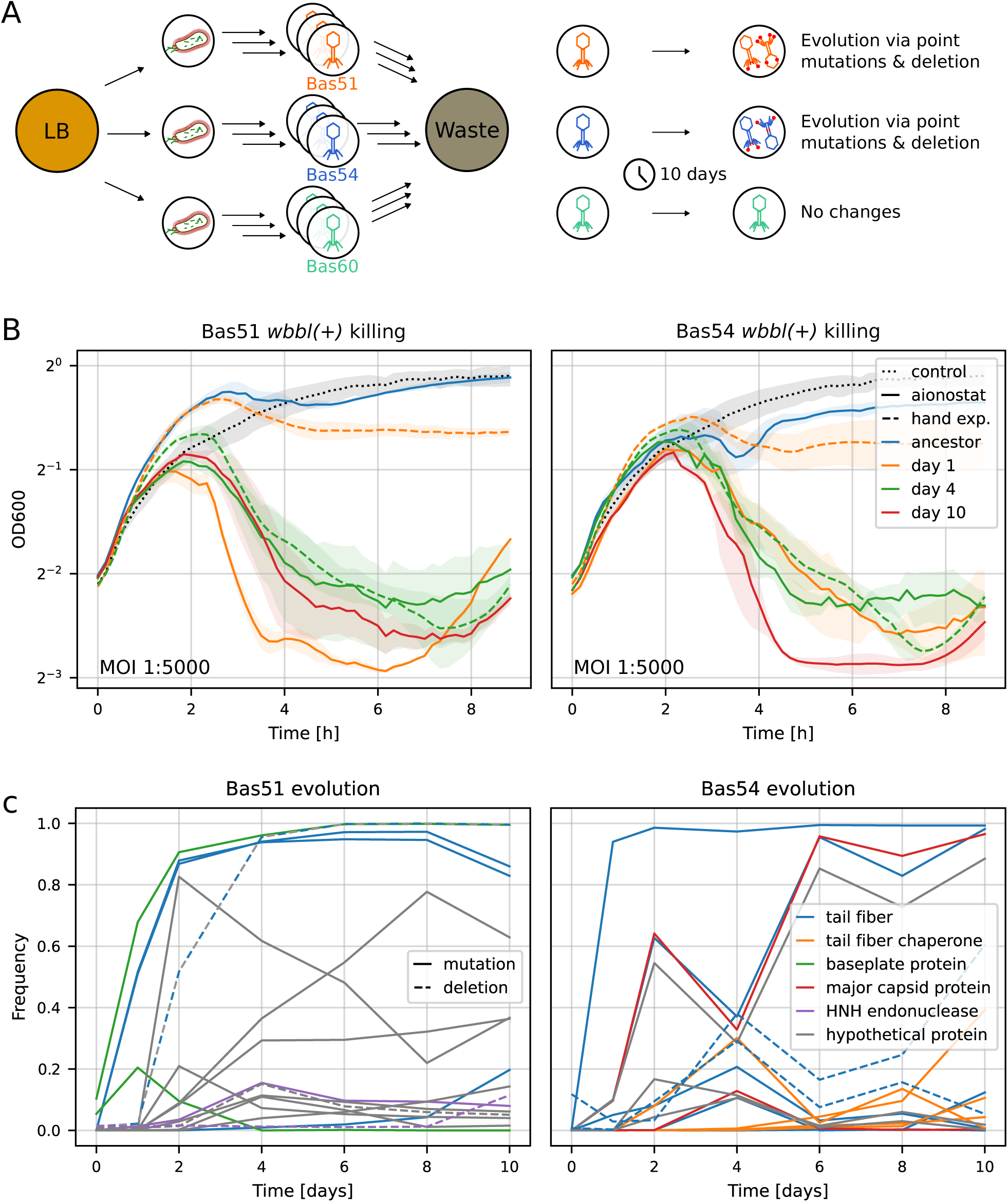
Linear evolution of phages. (**A**) Experiment schematic. Three phages were evolved in parallel, in separate vials and in triplicates, on *E.coli* BW25113 *wbbl(+)* for 10 days using the Aionostat. Samples of the phage populations were taken once a day. Evolution was observed for Bas51 and Bas54, which were analyzed in subfigure B and C. (**B**) Killing curves of ancestor phages and evolved populations on *E.coli* BW25113 *wbbl(+)* at different evolution time. Population evolved with the Aionostat achieve good killing after one day, while it takes 4 days with evolution using serial passages (labeled hand exp.). Evolved phages retain their killing efficiency on their isolation strain (see Supp. Fig. S3). (**C**) Frequency trajectories of genomic changes observed in the phage populations. Changes are mostly observed on genes related to host recognition and attachment. Point mutations are shown with solid lines, deletions as dashed lines. Colors represent the functional annotation of the genes where the changes occurred (where available).

**Figure 3:**
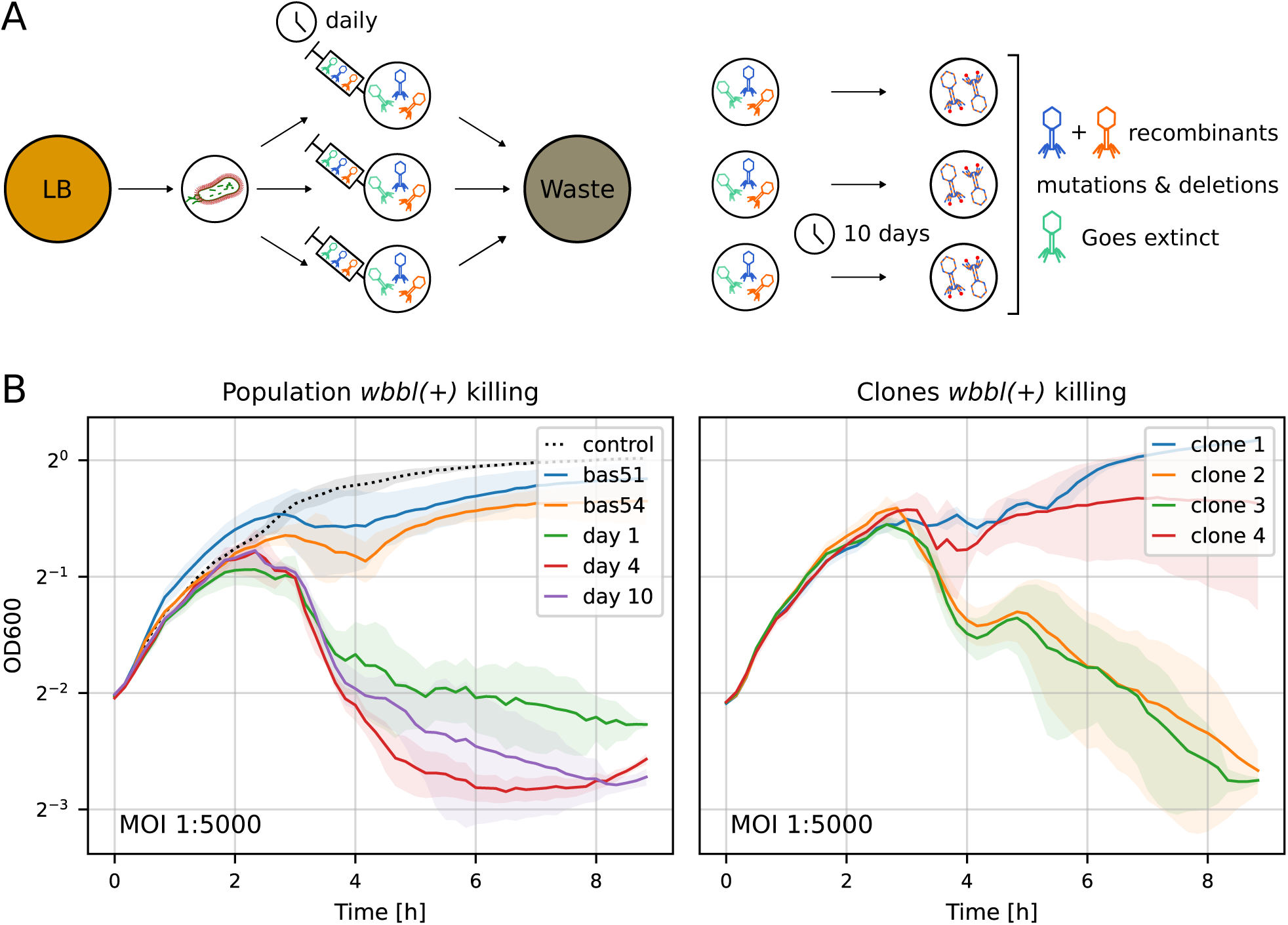
Evolution of phages via recombinations. (**A**): Experiment schematic. Three phages were mixed in equal amount and spread in 3 vials of the Aionostat. These cocktails were evolved on *E.coli* BW25113 *wbbl(+)* for 10 days. Samples of the phage populations were taken once a day before re-adding a small amount of the ancestral phage mix. **B**: Killing curves of ancestor phages, evolved populations and evolved phage clones (isolated from day 10 populations) from vial 1. The population of phages evolved via recombination show improved killing of *E.coli* BW25113 *wbbl(+)*. Isolated phage clones do not kill equally well, suggesting synergistic effects a the phage population level. The genome structure of these phage clones are shown in Figure 4.

Preparing, setting-up, and starting the experiment took about half a day, while around 20 min per day are required for maintenance and sample collection while the experiment is running. Samples taken from the feeder cultures were streaked on bacterial lawn, showing that the bacterial cultures stay phage free for the duration of the experiment. Phage concentrations in the phage vials were stable between 10^7^ and 10^10^ PFU/mL during the experiment, and were measured as explained in Material and Methods (M&M) section A.1. The ancestral phages, along with the phage populations from day 1, 2, 4, 6, 8 and 10 and phage clones from the day 10 populations, were sequenced using Oxford Nanopore technology as detailed in M&M A.6. Genomic changes were tracked over the experiment’s duration using the sequencing data, which also confirmed the absence of contaminations. Turbidity based killing curves were used to measure the improvement in phage fitness compared to reference phages. We present these results in the following sections, focusing on phage Bas51 and Bas54 as no significant evolutionary changes were observed for phage Bas60.

### Linear evolution of phages

#### Phage populations evolved with the Aionostat show improved killing

The linear evolution experiment using the Aionostat was performed as shown in schematic Figure 2.A. Figure 2.B shows the killing curves of *E.coli* BW25113 *wbbl(+)* by the ancestor and evolved phages starting from a multiplicity of infection (MOI) of 1 phage to 5000 bacteria. Details about the methodology can be found in M&M A.5. We can see in this figure that the ancestor phages Bas51 and Bas54 do not cause significant killing of the bacterial population in these conditions (blue line). The evolved phage populations cause a steeper decline in OD values than their ancestors between three and seven hours (other colors), indicating their improved killing efficiency on *wbbl(+)*.

Three randomly picked phage clones were isolated from plaques and amplified from each of the day 10 populations to measure their killing efficiency. Some phage clones showed limited killing of *wbbl(+)* similar to ancestral Bas51 and Bas54, while some others performed like the evolved phage population as a whole, see Supp. Figure S2. Evolved phage populations were also tested on the *E.coli* strain without O-antigen in Supp. Figure S3, showing conserved killing efficiency on this host. This was also the case for isolated phage clones, which shows their ability to infect both bacterial strains.

#### The Aionostat evolve phages faster than serial transfer evolution experiments

To compare the evolutionary dynamics in the Aionostat to manual serial passage, we evolved the same phage-bacteria combinations via daily passage and dilution, see section M&M A.2. The killing curves of the phages evolved with this approach are shown in Figure 2.B (dashed lines). These evolved phage populations originally show poor killing as highlighted at day 1 (dashed orange line). After four days of evolution (dashed green line) and later, the killing is similar to what we can observe with the Aionostat experiment. It seems that both approaches result in the same evolution outcome regarding killing efficiency, but the Aionostat allows for a faster evolution of phages. Interestingly, the improvement of killing seen between day 1 and 4 with the serial transfer approach coincides with the appearance of mutations at the same genes as in the Aionostat experiment for phage Bas51. Overall there are more genetic changes observed in the Aionostat experiments than in the serial transfer experiment. This suggests that the increased culture volumes and continuous replication in the Aionostat enable a more thorough exploration of the genetic landscape of beneficial mutations.

#### Genomic changes link improved killing to changes in host recognition genes

We sequenced the evolved phage populations at multiple time points to high depth. Using this population sequencing data, we determined the frequency trajectory of genomic changes (see Figure 2.C for changes that were seen above 10% frequency in at least one time point, M&M A.7 for details).

Figure 2.C shows how the originally clonal population of phages diversifies in the following days with diversity or selective sweeps at around 10 loci. The majority of changes were point mutations (full lines) with a few deletions (dashed lines) of varying length. Some mutations appeared and became widespread in the population in a matter of days, suggesting that they provide a strong fitness increase on *E.coli* BW25113 *wbbl(+)*. Notably, the deletion that fixed in the Bas51 population is a large 8.5kbp deletion that knocks out one of the tail fiber genes and following hypothetical proteins. We detected the same deletion in other replicates of the evolution experiment with this isolate. A deletion at a homologous locus was also observed at 5% frequency in Bas54 by day 10 (below the plot threshold), highlighting convergent evolution. We often observed mutations that rise before decreasing in frequency again, suggesting competition between mutations happening on different genetic backgrounds. Interestingly, we also saw that newly acquired mutations do not necessarily reach 100% frequency, keeping some diversity in the population. Unsurprisingly, most of the phage clones isolated from the day 10 population are genetically similar, and have the mutations seen at high frequency in Figure 2.C.

Comparing the timing of the emergence and spread of mutations to the killing curves in Figure 2.B, suggests that the earliest observed genomic changes are linked to the phages’ enhanced capability to kill the *wbbl(+)* strain. These mostly affect genes related to bacterial host recognition and attachment, such as the tail fibers or the baseplate protein. They likely impact the binding efficiency of the phages on *E.coli* BW25113 *wbbl(+)*. Adaptation of the attachment and entry process is also expected given that the bacterial strain used in the experiment is genetically identical to the isolation strain of the phages, except for the restored O-antigen. The O-antigen is thus the only new evolutionary challenge.

### Evolution of phages via recombination

#### Recombinant phages show improved killing

Next, we set out to test whether the Aionostat could be used to evolve phages via recombination. A schematic of the experiment and its results is shown in Figure 3.A. This experimental protocol is similar to the linear evolution experiment discussed previously, but the phage vials instead contained an equal amount mix of Bas51 Bas54 and Bas60 at the start. A small amount of this “cocktail” was added daily to maintain genetic diversity and provide opportunities for recombination during the entire experiment. In all vials, Bas60 disappeared, and the phage populations evolved by recombination of Bas51 and Bas54 in addition to point mutation and deletions.

Figure 3.B left panel shows the *wbbl(+)* strain killing curves from phages of the first vial. Evolved phage populations showed a rapid improvement in killing efficiency. The right panel shows the killing dynamics of the four clones isolated from the day 10 evolved population. Clone 2 and 3 show good killing efficiency, while clone 1 and 4 behave similarly to the ancestor phages. Comparing both panels suggests that using the phage population as a whole results in faster killing than the phage clones taken individually. Overall, these improvements in killing are similar to what we observed in the linear evolution experiment.

#### Recombinant phages dominate

To characterize the extent of recombination after 10 days of evolution in the Aionostat, we isolated four phage clones from each vial and sequenced their genomes. Figure 4 provides a graphical overview of these clones, highlighting the genomic segments inherited from Bas51 and Bas54.

**Figure 4:**
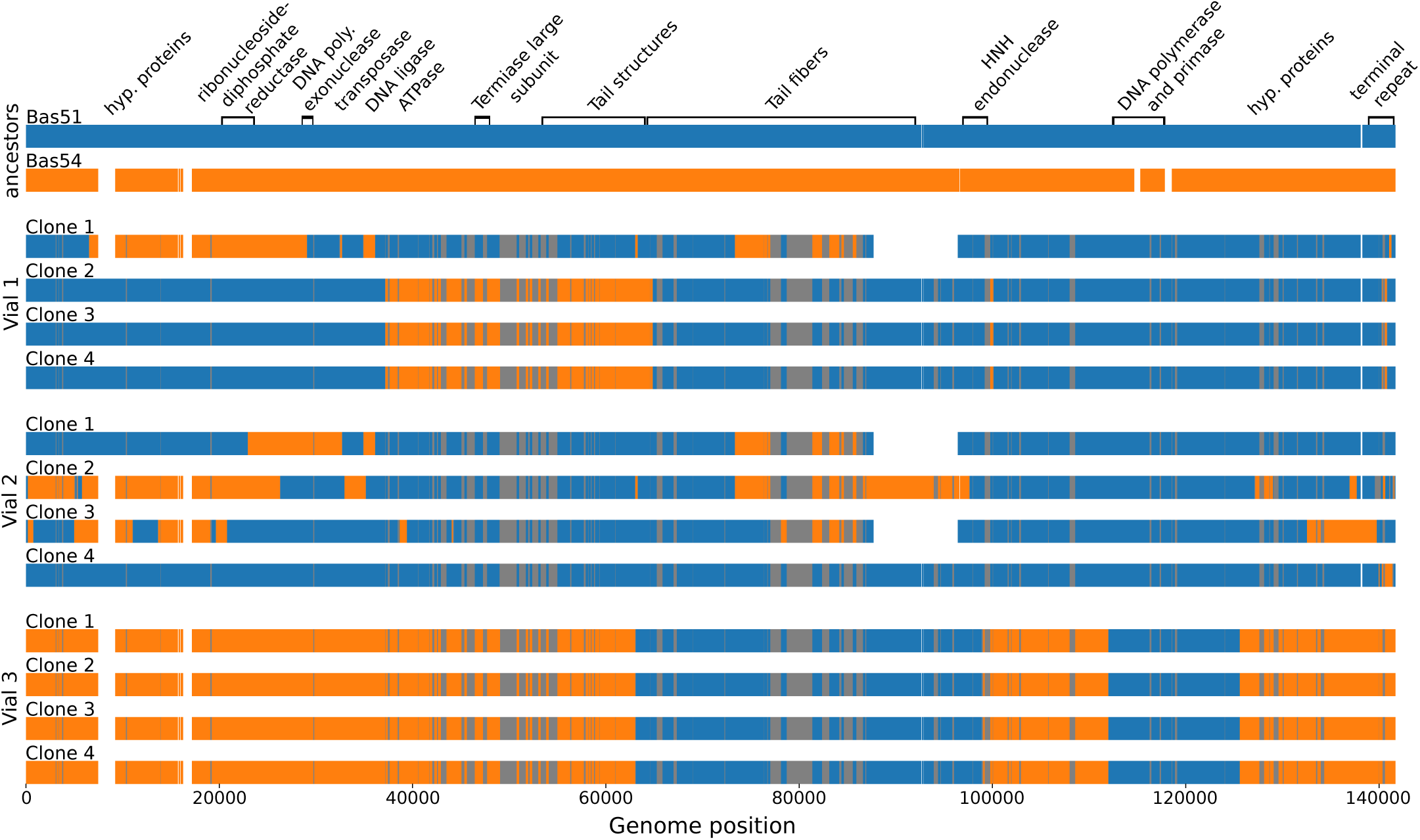
Genome alignment of recombinant phage clones. Four phage clones were isolated and sequenced from each of the Aionostat vials after 10 days of evolution as shown in Figure 3.A. Colors show the sequence similarity to the ancestor phages, grey is used when both are identical in a sequence stretch. White shows a missing genomic region. All phage clones are recombinants of Bas51 and Bas54, with some vials showing more diversity than others. No sequence stretch was acquired from Bas60. Some clones have large genomic deletions and point mutations relative to their ancestors. Reference sequences were rotated to improve visual clarity. See M&M A.11 for details of the methodology.

All clones are recombinants of Bas51 and Bas54, confirming that recombination is a common evolutionary route for phages (Hendrix, 2002; Hatfull, Hendrix, 2011). Notably, clones from vial 1 and 2 display diversity in genomic arrangements, while in vial 3 one particular recombination pattern is found in all four clones. The locations of the recombination breakpoints vary, but we observe similar patterns of recombination breakpoints near genes encoding tail fibers and in the region around an HNH endonuclease in different vials.

In addition to these mosaic patterns, several clones carry point mutations and large deletions near the end of the tail fiber locus, similar to the large deletion observed in the linear evolution experiment. Importantly, none of the clones contained any sequence stretch from Bas60, which had already disappeared from the culture. Altogether, these results demonstrate that the phage populations rapidly diversified via recombination and point mutations, creating composite phages adapted for improved infection of the *wbbl(+)* strain as shown in Figure 3.B.

#### Recombination dynamics at the population level

To quantify where in the genome recombination occurs, we analyzed single reads from the population sequencing data. Each read corresponds to a part of phage genome in the culture, and analyzing individuals reads allows us to generate a population summary of recombination. We built a Hidden Markov Model (HMM) to identify which stretch of each read originated from Bas51 or Bas54 and estimated the position of the recombination breakpoints. In Figure 5, the data from vial 1 is shown as the proportion of reads inferred as coming from Bas51 and Bas54, for each position, overlaid with the inferred breakpoints. Results for the two other vials are presented in Supplementary Figure S4 and S5. See M&M A.12 for details about the methodology.

**Figure 5:**
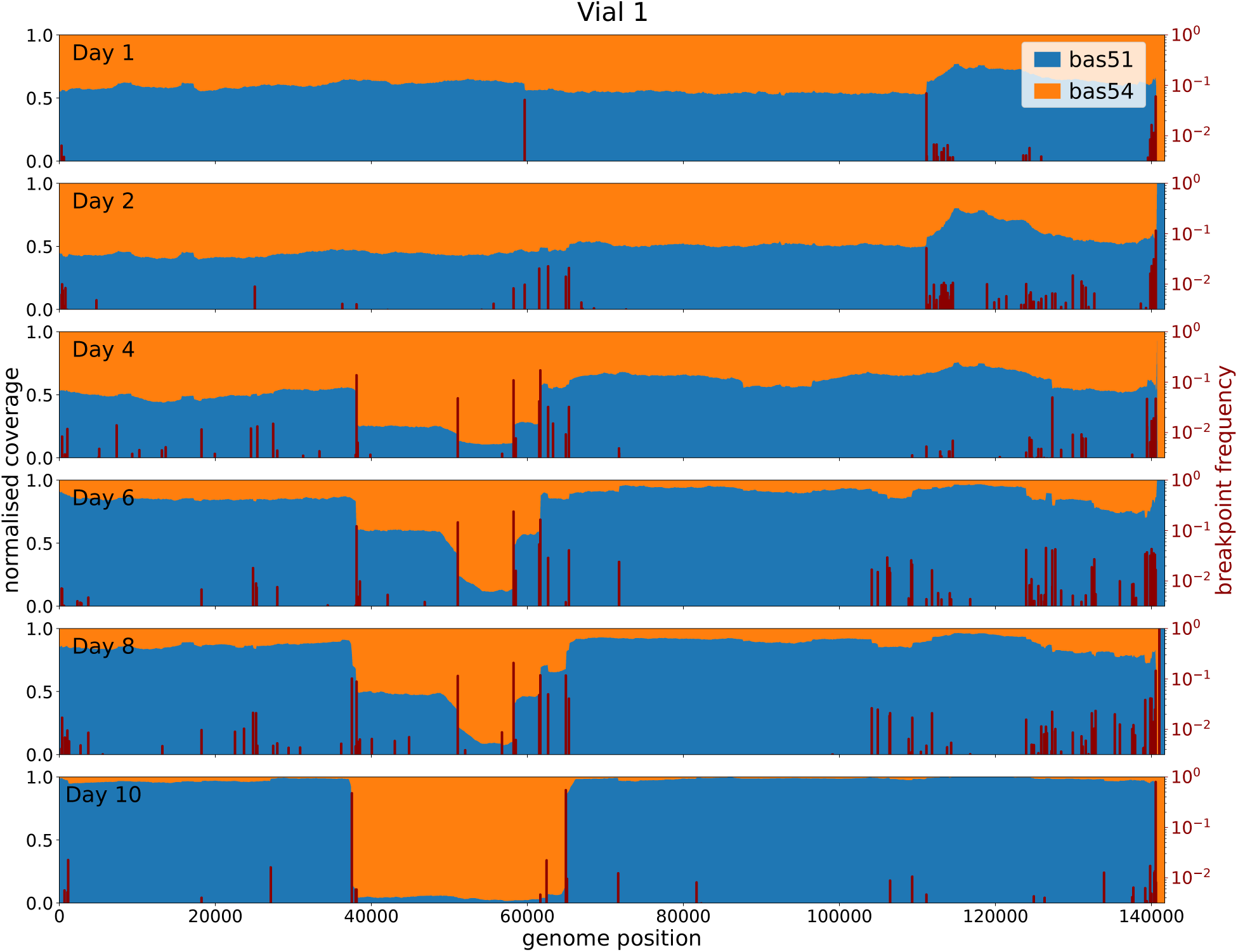
Recombination dynamics of evolved phage population. A Hidden Markov Model was used on the mapped population sequencing reads to determine which stretch of DNA originates from which ancestor at the single read level for all sequenced timepoints. This analysis allows us to show the proportion of each genome position coming from each ancestor phage in the population sample, which we report here as normalised coverage, alongside the recombination breakpoints inferred. At day 1 the population is mix of Bas51 and Bas54, while Bas60 has already disappeared. Over time we see the apparition of clear recombination breakpoints which correlate well with the change in normalised coverage. The final timepoint show that most phages from the population are from Bas51 origin, with a large recombinant part from Bas54. This corresponds to 3 of the 4 clones shown in Figure 4. See section M&M A.12 for methodology details.

At day 1, the population is a 50% mix of Bas51 and Bas54, with some clear bumps in the normalized coverage and inferred recombination breakpoints, indicating that recombinant phages started to appear. Many more recombination events were observed in vials at days 2 to 8, especially between genome positions 40,000 and 65,000, around 110,000, and from 125,000 to the end of the genome. By day 10, the diversity of the population had decreased, and most of the phage population are a Bas51 backbone with a large insert from Bas54 between position 37,000 to 65,000. This pattern corresponds to the genotype of three of the four clones shown in Figure 4.

Overall, these results show that recombination rapidly diversifies the phage population with hundreds of distinct recombination breakpoints at high enough frequency to be observed in population sequencing. The emergence of recombinant phages in the population likely contributed to the improved killing efficiency on *E.coli* BW25113 *wbbl(+)* observed in Figure 3.B.

## Discussion

Experimental evolution is a powerful tool for adapting phages to specific hosts and environments. But running such evolution experiments manually is labor-intensive and time-consuming. In this study, we introduced the Aionostat, a fully automated continuous culture device designed to facilitate high-throughput evolution experiments with phages. The Aionostat offers rapid, automated and reproducible evolution experiments, which are essential for understanding and leveraging phage evolution in various applications. Compared to existing continuous culture systems, the Aionostat offers enhanced versatility, reliability and accessibility. We demonstrated its capabilities to evolve phages in two distinct experiments: a linear evolution setup and a mixed “cocktail” experiment. The Aionostat generated phages with improved ability to infect and kill the target *E.coli* strain faster than manual serial passage experiments.

In the linear evolution experiment, the Aionostat rapidly produced phage populations with enhanced killing of *E.coli* BW25113 *wbbl(+)* through several point mutations and genomic deletions. These changes primarily occurred in host recognition and attachment genes, suggesting that improved adsorption was the key driver of adaptation to the restored O-antigen barrier. This observation aligns with recent work showing that adsorption is the key factor dictating *E.coli* phage host range (Humolli et al., 2024; Gaborieau et al., 2024). Similar mutations emerged across the three replicates and in a manually conducted serial-passaging experiment (although at a slower pace), indicating convergent evolution as a common adaptive response. A similar experiment showed that Bas51 Bas54 and Bas60 do not evolve when using *E.coli* BW25113, showing that evolution occurs to overcome the O-antigen barrier.

In the phage “cocktail” experiment, recombinant phages with improved killing of *wbbl(+)* emerged rapidly and displaced their non-recombinant counterparts, indicating a clear competitive advantage for recombinant phages. These phages are recombinants of Bas51 and Bas54, which are closely related reference phages, with only 2.3% mash distance (Ondov et al., 2016). These reference phages are lytic, which suggests recombination events happened when two distinct bacteriophages co-infected the same bacteria (Kunisaki, Tanji, 2010). No sequence stretches were acquired from the more distant Bas60 reference phage (∼ 40% mash distance). These phages exhibited diverse recombination breakpoints, and some vials displayed more genomic variety than others. By applying a Hidden Markov Model to the sequencing data, we could directly follow the dynamics of different recombination breakpoints at the population level. This was made possible by recent advances in Nanopore sequencing, which now provides reads long enough and of sufficient quality to capture large segments of individual phage genomes directly from the population.

While these experiments highlight the Aionostat’s capacity to drive rapid phage adaptation, they were primarily designed as a proof of concept rather than for deep biological inquiry. The choice of an *E.coli* with restored O-antigen as evolutionary challenge proved relatively easy for the phages evolved with the Aionostat, leading to robust killing within a single day. Further in-depth genetic analysis could be pursued to pinpoint the exact role the observed mutations in the phages’ improved predation capacities and confirm that it is linked to absorption efficiency on the bacteria.

Moreover, the 10 days run time of the experiment might have been excessive, since most of the adaptation in the phage populations was observed in the first few days. The evolution following the initial adaptation allowed bacterial populations to undergo phenotypic and occasional genotypic changes. Notably, in the phage vials, bacteria were also present that appeared to coexist with the phages, possibly through a mechanism of phenotypic heterogeneity (Wenner et al., 2023). Like any automated experiment device, the Aionostat does have limitations. Its complexity and the requirement for user calibration may challenge its user-friendliness, both in its construction and use. Additionally, its high throughput necessitates streamlined pipelines for sequencing and data analysis to keep up.

In conclusion, the continuous culture devices like the Aionostat promise to make to significantly accelerate phage research. It opens new avenues for understanding phage evolution at scale and provides a robust platform for other fields such as microbial evolution and therapy development.

## Acknowledgements

We gratefully acknowledge stimulating discussion with Reto Tschannen, Leonardo Lemos Rocha, Luca Sesta, Cornelius Roemer, Pablo Manfredi and Nicolas Wenner. This work was funded by the core funding of the University of Basel, SNSF grant 310030 188547, and the PhD Fellowhip Program of the Biozentrum (to VD).

## A Material and methods

### A.1 Measuring phage concentration

Phage concentrations were measured using serial dilution spotting on top agar. Large LB agar plates (12 cm x 12 cm) were overlayed with top agar (LB agar containing only 0.5% agar; stored at 60°C) supplemented with 300*µ*L of bacterial overnight culture. Subsequently, each 2.5*µ*L of all serial dilutions of phages were spotted on all top agar plates and dried into the top agar before incubation at 37°C for 4 hours. Plaques were counted from the lower dilution were single plaques were distinguishable, and this count was used to infer the phage concentration in the original solution in particle forming unit per milliliter (PFU/mL).

All phage concentrations were measured on both *E.coli* BW25113 and *E.coli* BW25113 *wbbl(+)* as we expected evolved phages could loose their infectivity on *E.coli* BW25113. This turned out not to be the case as both plates always showed similar concentration.

### A.2 Manual linear phage evolution experiment

This experiment is similar to the experiment done with the Aionostat presented in the main text but done by hand, to provide a comparison with a more standard approach to phage evolution experiments using daily serial passages. The experiment was performed with the same phages: Bas51, Bas54 and Bas60 in triplicates. Only the results of Bas51 and Bas54 are presented as not much evolution was observed for Bas60 (like in the Aionostat exepriment). The experiment lasted 5 days, as evolved phage population showed killing on *E.coli* BW25113 *wbbl(+)* similar to what was observed in the experiment performed with the Aionostat from day 4 samples.

On the first morning, 5mL of LB in glass tubes were seeded with 1*µ*L *E.coli* BW25113 *wbbl(+)* from an overnight culture and infected with the respective phages at an MOI of 1:100, allowing for several replication cycles before saturation in the vials. The cultures were then grown on a shaking incubator at 37°C for 8 hours, which was shown to be enough time for the culture to either lyse or saturate.

After 8 hours, 1mL samples were taken from each tube and cleared of bacteria using 1% chloroform plus strong vortexing, followed by a 2 minutes spin at 20 000g. The supernatant was extracted and stored at 4°C. It was used to seed the cultures on the next morning and later for the analysis of the phages at that time point.

The next morning, 10*µ*L of a 10^5^ dilution of the supernatant was used to infect the daily cultures prepared as was done on the first day. The amount of supernatant used ensured between 100 and 10 000 phages are transferred to the vial, preventing the extinction of the phages by over diluting while keeping the MOI low. These serial passages were repeated every day until the end of the experiment.

### A.3 Phage amplification

The samples taken from the evolution experiment usually contained between 10^7^ and 10^10^ PFU/mL. When more phages were needed, like in the case of DNA extraction for sequencing, the phage samples were amplified in liquid cultures. The amplification step was designed to minimize bias from the original sample by using an initially high amount of phages and a short incubation time to limit the number of replication rounds.

For each amplified phage stock, tubes were inoculated with 1mL LB and 300*µ*L of *E.coli* BW25113 *wbbl(+)* from overnight culture and then put for 20min at 37°C 600RPM to restart the growth of bacteria. Subsequently, 100*µ*L of the phage sample to amplify was added to the tubes and then incubated for 3 hours at 37°C 600RPM. The tubes were then cleared of bacteria by adding 1% chloroform, vortexing and spinning the tubes for 10min at 8000g. The surpernatant was extracted and titered as explained in M&M A.1, usually achieving between 10^10^ and 10^12^ PFU/mL. These amplified samples were stored at 4°C until use.

### A.4 Phage genomic DNA extraction

Genomic DNA of bacteriophages was prepared from high-titer stocks produced as explained in M&M A.3. The DNA was extracted using the Norgen Biotek Phage DNA Isolation Kit according to the manufacturer guidelines. When the DNA amount was to low for subsequent sequencing, the samples were concentrated using a SpeedVac vacuum concentrator. Quality of the DNA was controlled using a Nanodrop device and was sequenced as explained in M&M A.6.

### A.5 Plate reader killing curves

Killing curves presented in Figures 2, 3, S2 and S3 were generated using an Epoch2 plate reader in absorbance (OD600) mode. The phages were tested on both *E.coli* BW25113 and *E.coli* BW25113 *wbbl(+)* at a target multiplicity of infection of 1 to 5000. The curves shown are the mean of 3 replicates, with shaded area representing the standard deviation around that mean over the 3 replicates.

Each well was prepared with 180*µ*L of a bacterial dilution in LB of 5 · 10^8^CFU/mL. 20*µ*L of diluted phages with a concentration of 10^6^PFU/mL was then added to their respective wells, achieving a final phage concentration of 10^5^PFU/mL and a volume of 200*µ*L in the wells. The phage stocks were titered on the morning of the experiment to ensure as much accuracy as possible, and then diluted in PBS to hit the target MOI of 1 to 5000. The phage dilutions used to prepare the plate were also titered, as explained in M&M A.1, right after the plate was loaded to the plate reader to ensure the MOI was correct.

Once prepared, the plate was then moved to the Epoch2 plate reader and run for 15 hours at 37°C degrees, which was long enough to observe bacteria killing and eventual regrowth of resistant bacteria. The experiment was performed with 450RMP double orbital rotation and OD600 readings every 10 minutes. The lid of the plate was removed, and replaced with a Breathe-Easy sealing membrane.

### A.6 DNA sequencing

The DNA samples extracted as explained in M&M A.4 were sequenced in-house using the Oxford Nanopore sequencing technology. We utilized the MinION Mk1B device for sequencing, employing V14 chemistry coupled with R10.4.1 pores. The flow cells used in this procedure were of the type FLO-MIN114. To facilitate the sequencing, we utilized the rapid barcoding sequencing kit 24, specifically the kit SQK-RBK114.24. For the basecalling process, Dorado version 0.7.0 was employed, using the basecalling model dna r10.4.1 e8.2 400bps sup version 5.0.0. The basecalling pipeline used is available here: https://github.com/vdruelle/nanopore_ basecalling.

### A.7 Mutation trajectories analysis

The analysis of the mutations shown in Figure 2.C was performed using a Snakemake pipeline, which is publicly available at https://github.com/mmolari/evo-genome-analysis. This pipeline takes as input the raw reads from the samples and the reference genomes, obtained as explained in M&M A.6. It maps the reads from each sample to the references using Minimap2 (Li, 2018) from which we extract trajectories of genomic changes over time, encompassing single nucleotide polymorphisms, gaps, insertions, clips, and rearrangements. These trajectories were then filtered and the ones seen above 10% in at least one time point were plotted as shown in Figure 2.C. These mutations were then manually inspected and categorized based on their annotation.

### A.8 Bacteriophage reference sequences

The reference sequences used in the analysis of recombinant phages are rotated with respect to those present in NCBI GenBank (Bas51: MZ501111.1; Bas54: MZ501093.1). The assemblies deposited in the database were created with short-read sequencing technology (Illumina) and contain the terminal repeat region (the region of repeated motifs representing the molecular termination of the DNA molecule) in the middle of the genome (Bas51: 93,911–96,377; Bas54: 92,785–95,298).

The genome assembly procedure used in this study detailed in M&M A.9 is based on long-read sequencing described in M&M A.6, which enables the unambiguous determination of the relative positions of the sequence components. Not having the terminal repeats at the end of the genomes could generate artifacts that could be interpreted as a recombination breakpoint if the start and end of the phage genome originates from different ancestor phages. Therefore, to increase visual and biological consistency, the two GenBank reference sequences were rotated so that the terminal repeat region appeared at the end (starting at 96,377 and 95,298 for Bas51 and Bas54, respectively).

### A.9 Genome assembly

The recombinant phage clones were assembled with Flye 2.9.2 (Kolmogorov et al., 2019). The parameters used for the phage genome assembly were:

- `--nano-hq`, to handle ONT reads basecalled as described in M&M A.6
- `--genome-size` set to the length of the GenBank references (approximately 140 kb)
- `--asm-coverage` set to 40X, specifying the target coverage for the initial disjoint assembly

### A.10 Hybrid reference

For the analyzes presented in Figures 4 and 5, all sequences were aligned to a hybrid reference to keep the same frame of reference. The hybrid reference is an artificial sequence constructed from the multiple sequence alignment (MSA) of the two reference sequences from Bas51 and Bas54 that were rotated as explained in M&M A.8. In regions where both references map, the hybrid reference inherits a nucleotide from either sequence randomly. In genomic regions present in only one of the references, the hybrid reference takes the information from the sequence that is present.

### A.11 Genome alignment of recombinant phage clones

The individual lines from Figure 4 are generated by mapping the ancestral phage sequences onto the recombinant clone sequence assembled as explained in M&M A.9 and M&M A.10. From the alignment, the distribution of differences along the genome (mismatches and gaps) for each ancestral sequence is extracted. These difference counts are convoluted with a 250bp window and normalized by the difference distribution between the two ancestral sequences and the clone.

In regions where the normalized score of differences is less than 40% for one of the ancestral references, the clone genome is assigned to that reference (therefore colored with the color of the corresponding ancestral reference). On the other hand, when the score of differences is above that threshold for both references, the clone genome region is colored in gray. These regions correspond to the borders of recombination events, where the convolution window blurs the boundary, or to stretches in which the ancestral sequences are identical and it is impossible to determine the origin of the clone.

The convolution size and threshold values were chosen to sum mismatches over a sufficiently large region so that short identical stretches of nucleotides do not cause interruptions in the mismatch line. At the same time, the window is not too large, preventing overlaps between the mismatch distributions of the two references that would enlarge the gray regions at the recombination boundaries.

The script generating the plot can be found at: https://github.com/kcajj/recombinant_phages_study

### A.12 Hidden Markov Model (HMM) inference

Figure 5 is generated using a Hidden Markov Model (HMM) to infer the most likely path of each read through the two ancestral sequences. This inference is challenging due to the presence of noise in the ONT reads and the low density of informative sites between Bas51 and Bas54 (the genomes are about 97% identical). The reads coming from the recombinant populations are aligned to the hybrid reference (M&M A.10) and the differences with respect to the two ancestral sequences are extracted and used to create the HMM as explained below and illustrated in Figure 6.

**Figure 6:**
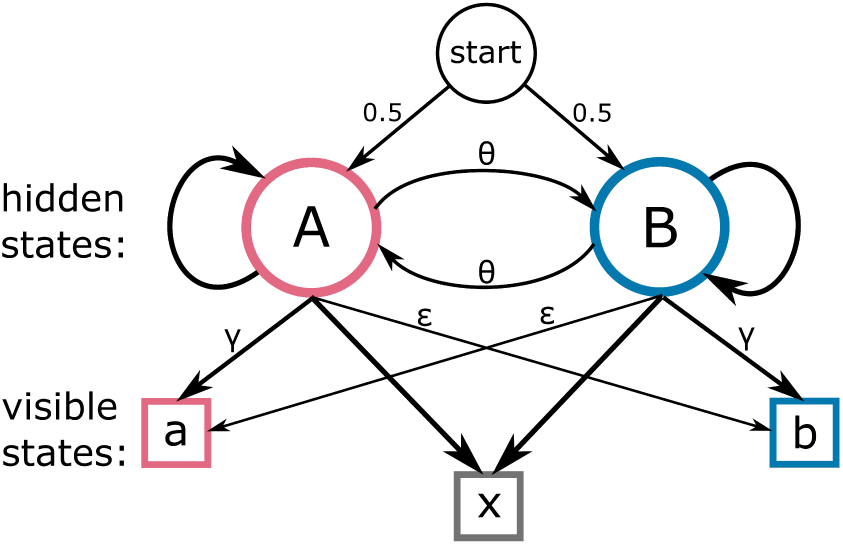
HMM model schematic. The two hidden states (A and B) correspond to the two reference phage genomes from which each read has been generated. The visible states (a, b and x) correspond to the evidences that are observed when the read is aligned to the two references.

The HMM has three visible states (evidence of the clone being similar to phage Bas51, Bas54 or no discernible evidence), emitted by two hidden states (corresponding to the two ancestral references). The probability of emitting each visible state depends on the hidden state. In the emission matrix, *γ* is the probability that a hidden state emits a correct evidence (i.e., ancestral A emits *a*, or ancestral B emits *b*), and *ɛ* is the small probability of emitting a misleading evidence (i.e., A emits *b* or B emits *a*). Since the two references are identical at most sites, there will often be no discernible evidence (denoted as *x*).

The probability *γ* (correct evidence) represents the fraction of the genome that differs between the two references minus the error rate of the sequencing technology. The probability *ɛ* (misleading evidence) corresponds to random sequencing errors suggesting the base of the wrong ancestral phage. The transition probabilities between hidden states are governed by a transition matrix, where *θ* is a hyperparameter related to the recombination frequency in the dataset. The model uses the Viterbi algorithm to find the optimal sequence of hidden states from the observed sequence of emissions. The emission parameters of the model were estimated empirically, measuring the frequencies of each visible state in a dataset with a known hidden state (separate sequencing runs of Bas51 and Bas54). The assumption behind such estimation is that long reads coming from a pure sequencing run of a phage present the same characteristics as the portions of reads of a recombinant phage population originating from the same phage. The parameters are:

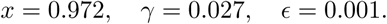

The value of the transition parameter *θ* was optimized by running the HMM on a subset of the recombinant population dataset across different values of *θ* and selecting the inference with the highest log likelihood. A peak in the total log likelihood was observed at approximately *θ* = 4 · 10*^−^*^5^. The highest log likelihood corresponds to the model that more closely fits the data. This value correlates with the recombination frequency in the dataset, suggesting an expected recombination event roughly once every 25kbp.

The accuracy of the model was tested on artificially generated sequences. An accuracy of 0.9986 was achieved in correctly assigning hidden states. Although breakpoint positions are consistent, they can not be inferred exactly in sequence streches that are identical between Bas51 and Bas54.

In Figure 5, for each position of the hybrid reference (x axis, M&M A.10), the proportion of reads mapping in that position inferred by the HMM as originating from Bas51 and Bas54 is represented by the proportion of blue and orange (left y axis). On top of this, the breakpoint frequency is represented. That is, the proportion of reads that present a breakpoint in that specific position according to the HMM inference (right y axis).

The pipeline used to analyze the recombinant population reads can be found at: https://github.com/kcajj/recombinant_population_analysis

### A.13 Aionostat components

This study showcased the ability of the Aionostat to perform directed evolution experiment on bacteriophages to improve fitness and killing on a challenging bacterial strain. In this section, we dive into the details of the Aionostat to present how it is able to perform such evolution experiments.

The Aionostat is an autonomous continuous culture machine that has two main parts. The first is composed of the components that handle the liquids, the structure for such components as well as the pumps that move the liquids around. This part sits inside an incubator for temperature control. The second part act as the brain and power source for the machine, which can be programmed to perform a wide variety of experiments. Overall, the Aionostat was made from commercially available components, as well as custom 3D printed parts and electric wiring.

#### Vials

The experimental setup utilizes vials of two sizes, specifically 8 mL and 40 mL total volume, as depicted in Supp. Figures S1.B and S1.C. These vials are interchangeable depending on the experimental requirements. For the experiments showcased, the larger vials were employed for both the bacterial and phage cultures. Alternatively, the smaller vials offer an option to intensify selective pressure on the phages. Utilizing a reduced volume for the phage culture effectively increases the dilution rate, which enhance selective pressure. This increased pressure can accelerates the sweeping of beneficial mutations, potentially leading to more rapid phage evolution.

Each vial is equipped with a magnetic stir bar to ensure consistent mixing. The vials are sealed with an open cap, fitted with a PTFE-coated silicon septum. This design allows for the sterile transfer of liquids in and out of the vials using needles and tubing. These components are shown in Supp. Figure S1.A.

#### Vial holders and magnetic stirrer plate

The vials are placed within custom 3D-printed holders. These holders not only secure the vials but also position the OD and level sensing sensors in close proximity to the vials, as illustrated in Supp. Figure S1.E. The vial holders, along with their respective vials, are positioned on a 15-spot magnetic multistirrer. They are held in place using acrylic panels crafted through laser cutting and are assembled with screws and 3D-printed spacers, as depicted in Supp. Figure S1.I.

#### Liquid handling

Liquid transfer into and out of the vials is facilitated by commercially available needles, which go through the silicon septum. These needles are connected on one end to silicon tubing using Luer connectors, and to piezoelectric pumps on the other end. These pumps offer more control over the flow of liquid in each tube and are more compact than single channel peristaltic pumps. This gives great versatility to the experiments that can be performed with the Aionostat. The pumps are organized in custom 3D-printed arrays, available in various configurations (5, 8, or 15 pumps), ensuring stable positioning of the pumps and separation of electrical connections from the tubing. The 5 pumps version is shown in Supp. Figure S1.G and S1.H.

Liquid from the input solution is pumped from the sterile bottles sitting outside of the incubator using pass through caps connected to the tubing and the piezoelectric pumps (Supp. Figure S1.F). Additionally, a 15-channel peristaltic pump is used as exhaust and overflow protection for the vials. The depth of the needle attached to the peristaltic pump is what sets the working volume in the vials. We used 60mm needles, which set the working volume of the vials to half of their total volume. This pump sits outside of the incubator. The flow rate of the piezoelectric and peristaltic pumps is calibrated before each experiment as explained in A.14.

#### Optical density measurement

Bacterial density within the vials is assessed using an optical setup involving an LED and a phototransistor positioned on opposite sides of the vial at a 135° angle. This configuration allows for the detection of light diffracted by bacteria within the vial, rather than relying on direct absorption, thus enhancing sensitivity at low bacterial densities. To accurately translate the phototransistor’s signal into optical density values, calibration against standards with known optical densities is essential. The detailed calibration procedure is outlined in the M&M A.14.

The LED and phototransistor are positioned using designated holes on the sides of the vial holders as shown in Supp. Figure S1.E. The specific models used are the MT5880-IR LED from Marktech Optoelectronics and the SFH 300 FA-3/4 phototransistor from ams OSRAM (Supp. Figure S1.D). Several combination of LEDs and phototransistors were tested, this particular pairing was found to offer the best dynamic range and signal-to-noise ratio for our experiments.

#### Liquid level sensing

To enhance control and safety in liquid handling within the Aionostat, custom capacitive sensors have been developed for monitoring the liquid level in the vials. These sensors are positioned in the inner section of the vial holders, ensuring direct contact with the vials. Given the fixed diameter of the vials, the volume of liquid present can be accurately determined from the readings provided by these sensors. In the experiments conducted, a constant volume was maintained, so we used these sensors primarily as a means of overflow protection. While beneficial for safety, these sensors are not essential for conducting such experiments at a constant volume.

#### The controller

The Aionostat is controlled using electronic components that are outside of the incubator and connected to the components inside via wires. Its central processing unit is a Raspberry Pi 4B board, equipped with HATs for analog-to-digital conversion of voltage readings and additional GPIOs. Custom Python scripts were developed to automate the experimental procedures and are accessible at https://github.com/vdruelle/Aionostat_code. The analog to digital readings are used to measure the sensors signal, which provides feedback on the state of the experiment. The GPIOs control the activation of piezoelectric and peristaltic pumps, based on the feedback from the sensors and the ongoing experimental requirements. The pumps have a near constant flow rate, so the volume pumped is controlled by the time the pump is running.

### A.14 Aionostat protocols

#### Initial tests and calibrations

Calibration of the Aionostat’s components is a critical pre-experimental step. Firstly, the optical density sensors are calibrated. To perform the calibration, vials are prepared with bacterial dilutions of different optical density measured externally on a spectrophotometer. We also include a vial with raw media to cover optical densities between 0 and 1. Each vial holder is tested sequentially with these OD standards, recording the phototransistor’s voltage output. A linear fit between these voltages and OD values is computed and saved for each vial holder. During experiments, these fits are used to deduce the OD from the sensor signals. Calibration of each vial enables to compensate for component to component variability.

We continue with the calibration of the piezoelectric and peristaltic pumps. Given their constant flow rate, calibration involves running the pumps for a set duration and measuring the output volume. The flow rate is determined by dividing the volume by the time. These rates also enable calculation of the dilution rates in the vials by factoring the working volume in the vials.

Finally, if the level sensors are utilized, they also require calibration. This process involves recording sensor voltages with vials at varying liquid levels: empty, full, and intermediate volumes. The sensor readings and level of the liquid have a linear relationship, enabling the interpolation of liquid height through a linear fit.

#### Sterilisation of the Aionostat

Sterilization of the Aionostat is conducted post-calibration and pre-experiment to prevent contamination. This is achieved in two stages. First, all tubing, pass-through caps, and vials are washed and then autoclaved at 120°C for 20 minutes. The vials are assembled and sealed as depicted in Supp. Figures S1.B and S1.C, while the tubing and caps are wrapped in aluminum foil prior to autoclaving. These components are then installed in the Aionostat, and the tubing, needles, and pumps are connected as they will be during the experiment.

The second stage involves chemical sterilization using 3% sodium hypochlorite (bleach) and 3% citric acid solutions, applied sequentially throughout the entire setup with sterile MiliQ water used to rince inbetween and at the end. Beginning with the sodium hypochlorite solution, all pumps are activated for one minute, ten times sequentially over a 30-minute period. The vials are then emptied using the pumps, and this procedure is repeated with sterile MiliQ water and then the citric acid solution. After this, the vials are emptied again, and the system is rinsed with MiliQ water, which is run through the vials and tubing. This process is automated, requiring manual input only for changing the input bottles. At this point the inside of the Aionostat is sterile, and it is crucial to not disconnect any tubing.

#### Setting up

The next step is programming the Aionostat for the experiment, which involves using and modifying the pre-written control code. This can also be performed before or during the previous steps.

After programming and ensuring the Aionostat is set up and sterile, the MiliQ water in the vials is replaced with fresh sterile media. This replacement is carried out by changing the input bottle (near a flame for sterility) with fresh sterile media and using the pumps to exchange the liquid in the vials. Lysogenic broth was used in the experiments presented.

The final preparation step is the inoculation of the sterile media with bacteria and bacteriophages. This is done manually using a syringe and needle to pierce the septum and introduce the appropriate bacterial strain or bacteriophage into each vial. We finish by closing the incubator and setting it to the desired temperature. Typically, the entire preparation of the Aionostat takes 4 to 8 hours.

#### Running the experiment

At this stage the experiment is fully prepared, and we start the run by launching the code for the experiment from the Rasberry Pi. During the subsequent days, routine maintenance involves changing the input media bottles before depletion and replacing the waste bottle when it is full. A 5-liter bottle pre-filled with disinfectant, is used as the waste container. Manual sampling from the vials is conducted using a syringe and needle, piercing through the septum for sample collection. For storage purposes, bacteria were removed from phage samples. Upon completion of the experiment, the vials are emptied and disassembled, and the setup can prepared for the next experiment following the previously outlined steps.

Although no contamination was observed during the experiments presented in this study, previous trials have shown two instances of contamination. The first was a phage vial contaminated by another phage sample during the initial setup. The second was backgrowth of the bacteria in the input media bottle. This happened when the bottles ran out of media for a short time before being changed. We assume it allowed the bacteria to grow in the tubing in the absence of flow.

## A Supplementary materials

**Supp. Fig. S 1:**
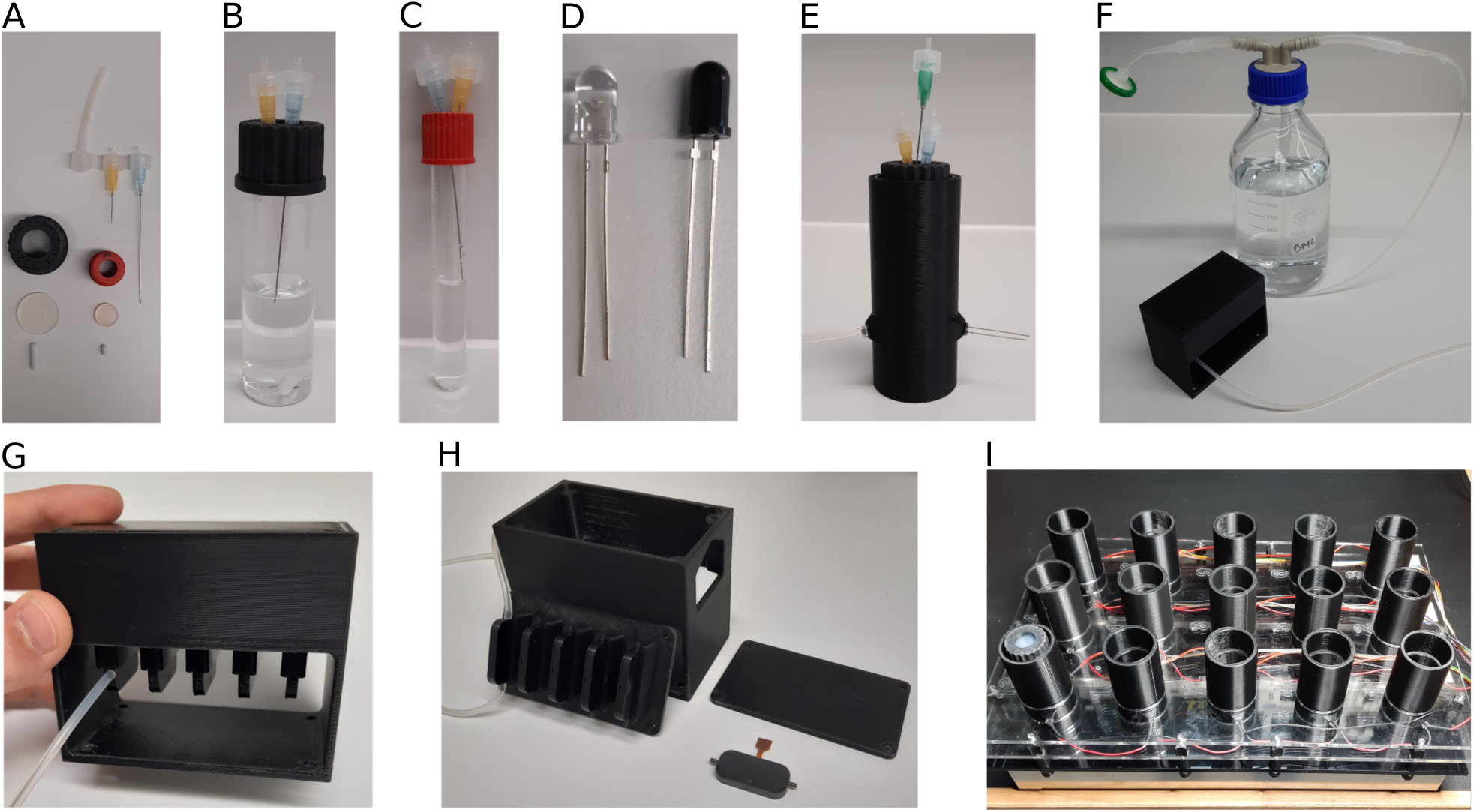
Components of the Aionostat. **A**: Needles, silicon tubing, Luer connector, open vial caps, silicon septum and magnetic stir bars used in the assembly of the vials. **B**: Big vial assembly (40mL). **C**: Small vial assembly (8mL). **D**: LED and phototransistor for the bacterial density measurement. **E**: Full assembly of one vial in its vial holder. This assembly is what is shown on the schematic in Figure 1.A. **F**: Pump connection to the input bottle. **G-H**: Piezo electric pump array and housing (5 pumps version). **I**: Vial holders positioned on the magnetic stirring plate. These components are combined to form the Aionostat as shown in Figure 1.B.

**Supp. Table. S 1:** Details about the genomic changes shown in Figure 2.C. The mutations are ordered by maximum frequency observed at any timepoint (from top to bottom) for each ancestor phage. The positions in bold are mutations that are also observed in the other two replicates of each phage population, showing signs of convergent evolution. Mutations are often observed on the same target genes in other replicates, even though not always at the exact same position.

| Phage | Position | Mutation | Gene |
| --- | --- | --- | --- |
| Bas51 | **43283–51785** | Deletion | Tail fiber, Hyp. prot, Head maturation protease |
|  | **33523** | SNP | Baseplate protein |
|  | **28953** | SNP | Lateral tail fiber with fibronectin type III domain |
|  | **30442** | SNP | Lateral tail fiber with fibronectin type III domain |
|  | **8536** | SNP | Hypothetical protein |
|  | 8030 | SNP | Hypothetical protein |
|  | 80615 | SNP | Hypothetical protein |
|  | 87618 | SNP | Hypothetical protein |
|  | **33761** | SNP | Baseplate protein |
|  | 39040 | SNP | Short tail fiber protein |
|  | 114994 | SNP | Putative homing endonuclease |
|  | 108939–109049 | Deletion | ClpP-like protease |
|  | 79997 | SNP | Hypothetical protein |
|  | 99662 | SNP | Hypothetical protein |
|  | 111490–113528 | Deletion | Reductase activator, Endonuclease |
|  | 106269 | SNP | Hypothetical protein |
| Bas54 | **36619** | SNP | Lateral tail fiber with glycosidase and deacetylase domains |
|  | 27128 | SNP | Tail fiber protein |
|  | 7207 | SNP | Major capsid protein |
|  | 98288 | SNP | Hyp. prot |
|  | **47397** | Deletion | Lateral tail fiber with intimin domain |
|  | 38314 | SNP | Tail fiber chaperone |
|  | **38346** | SNP | Tail fiber chaperone |
|  | 45228–45234 | Deletion | Lateral tail fiber with intimin domain |
|  | 31914 | SNP | Lateral tail fiber with fibronectin type III domain |
|  | 8516 | SNP | Hyp. prot |
|  | **7695** | SNP | Major capsid protein |
|  | 47464 | SNP | Lateral tail fiber with intimin domain |
|  | 31648 | SNP | Lateral tail fiber with fibronectin type III domain |
|  | 38337 | SNP | Tail fiber chaperone |
|  | 8537 | SNP | Hyp. prot |
|  | **38316** | SNP | Tail fiber chaperone |

**Supp. Fig. S 2:**
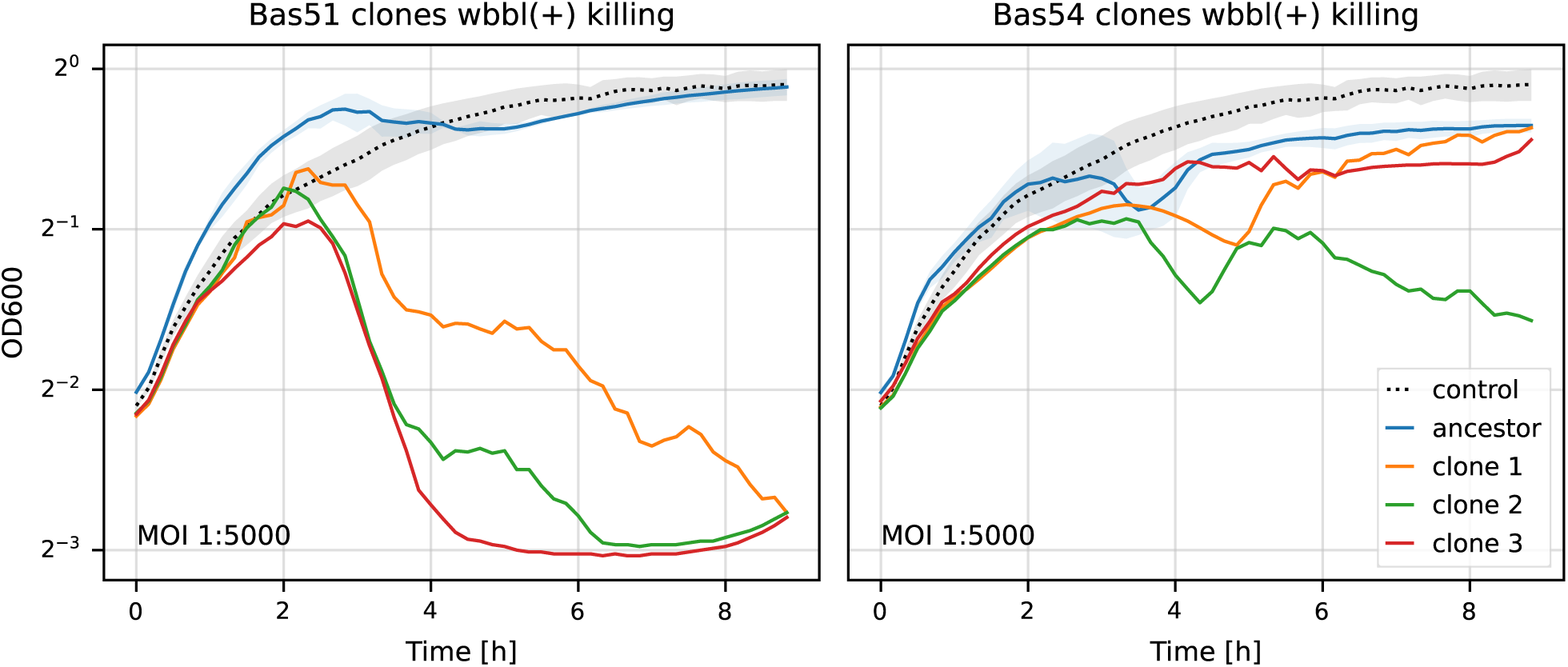
Killing curves of phage clones from linear evolution experiment. Clones were isolated from day 10 evolved populations. The blue lines are the same one as shown in Figure 2.B. The killing of the *wbbl(+)* strain is better than the ancestor phages for all clones, but clones from Bas54 seem only marginaly better than their ancestor. Phage populations as a whole seem better at killing overall (as seen in Figure 2.B).

**Supp. Fig. S 3:**
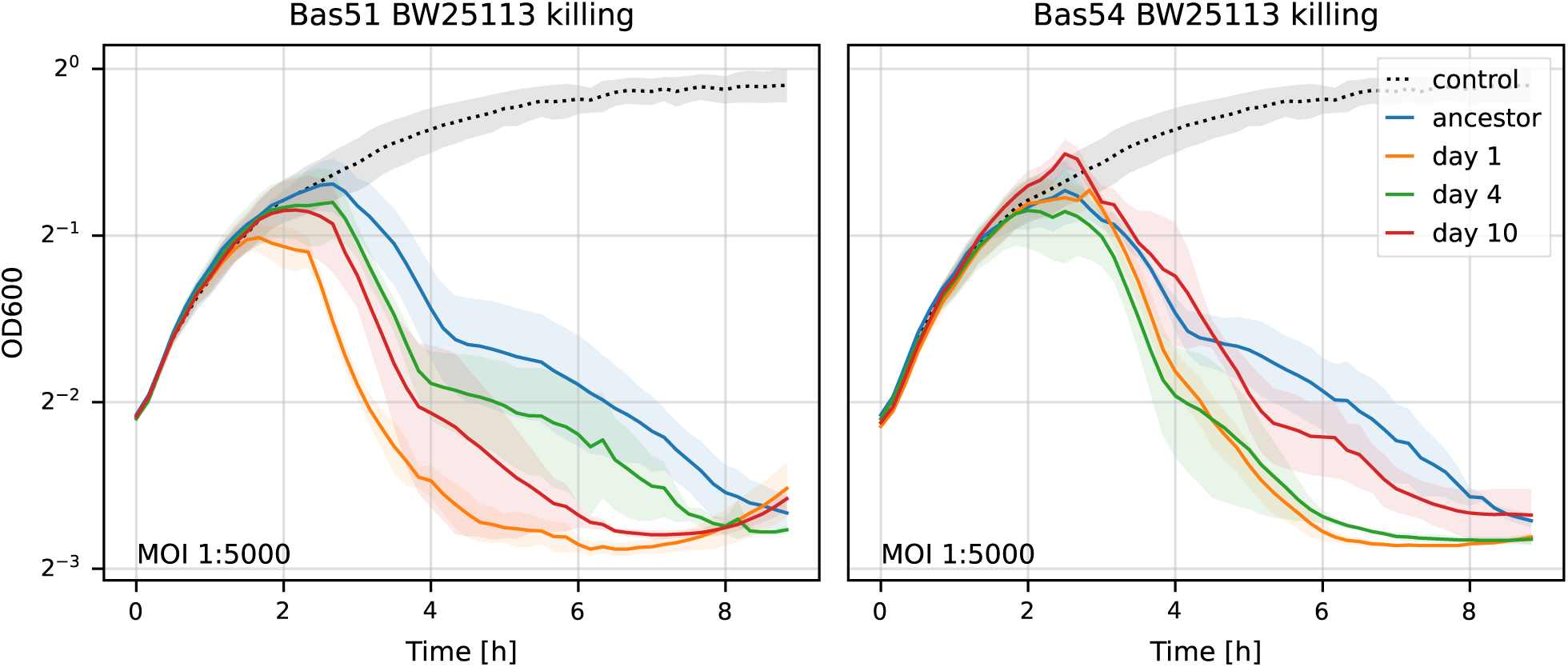
Killing curves of *E.coli* BW25113. This is the same as Figure 2.B but done on *E.coli* BW25113 rather than *E.coli* K12 *wbbl(+)*. The evolved populations seem to retain their killing efficiency on this relative of their isolation strain.

**Supp. Fig. S 4:**
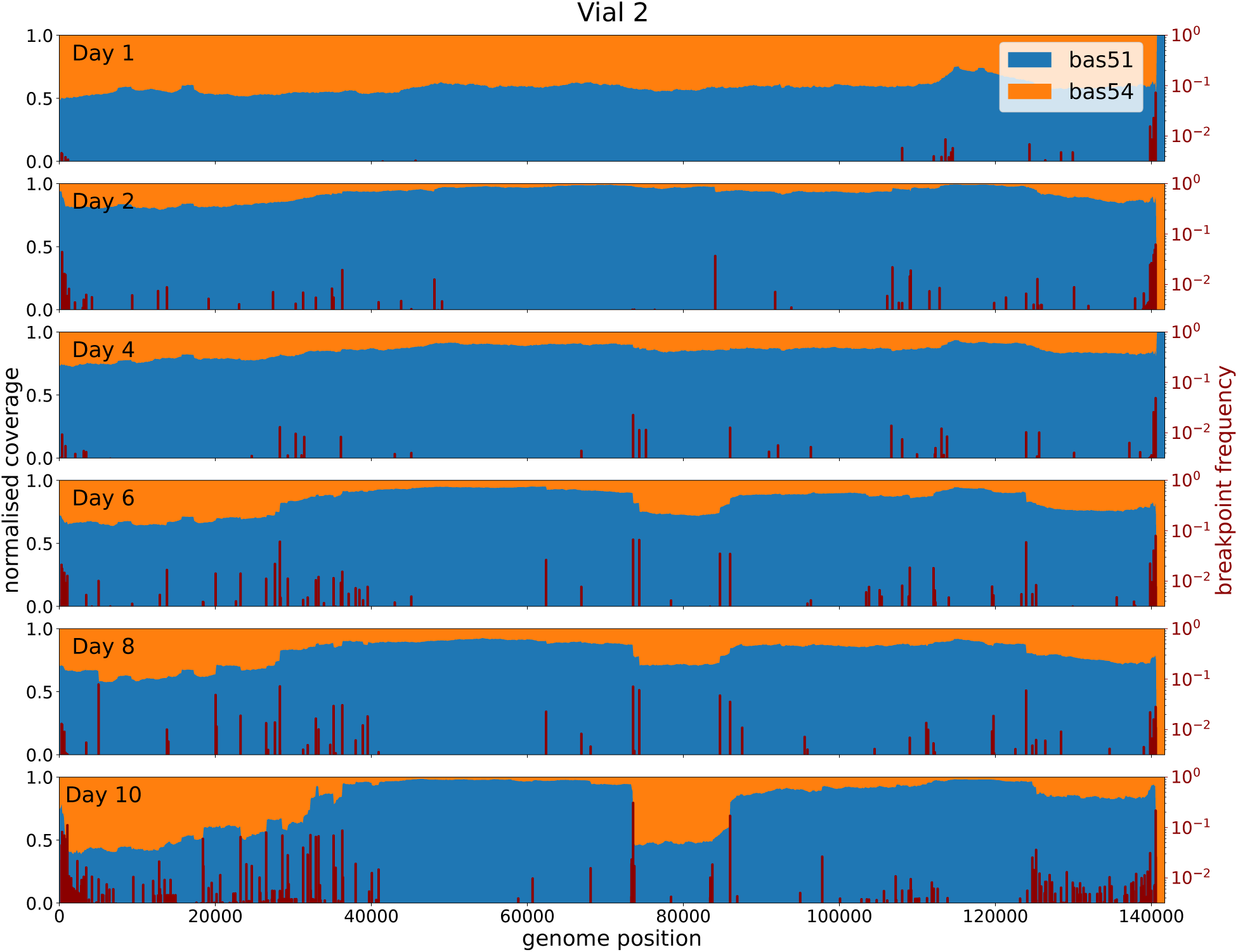
Recombination dynamics of evolved phage population from vial 2. This plot is similar to Figure 5 but from the data of the second vial of the recombination experiment instead.

**Supp. Fig. S 5:**
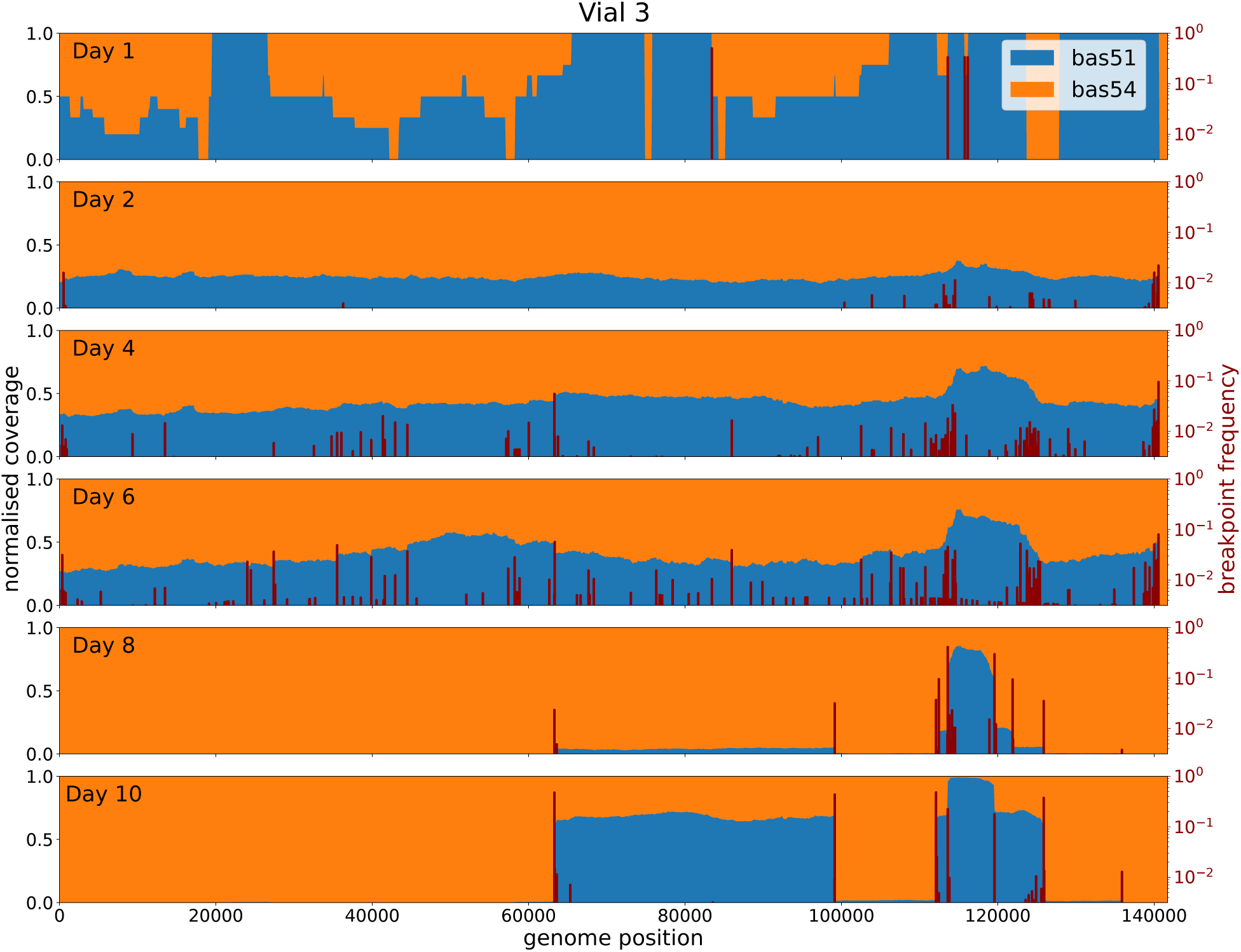
Recombination dynamics of evolved phage population from vial 3. This plot is similar to Figure 5 but from the data of the third vial of the recombination experiment instead. Interestingly this is the only vial where we saw a significant presence of Bas60 in the population at day 1. This explains the low resolution of the day 1 subplot (which only accounts for Bas51 and Bas54). Bas60 then disappeared in the following days as recombinants of Bas51 and Bas54 outcompeted it as seen in the other vials.

